# In Silico Identification of lncRNAs in Wheat Leaf Rust Pathogen Interactions

**DOI:** 10.64898/2026.11.19.767801

**Authors:** Barri Yaswitha, Jyothsna S, Alagu Manickavelu

## Abstract

Wheat (*Triticum aestivum* L.) is a major crop grown on about 225 million hectares worldwide, and it suffers heavy losses from rusts caused by fungal pathogens. This study focuses on leaf rust, caused by *Puccinia triticina*. When wheat is attacked by this fungus, hundreds of downstream genes in various biological pathways show regulated expression, which depends partly on non-coding RNAs (ncRNAs), including microRNAs (miRNAs) and long non-coding RNAs (lncRNAs). Because lncRNAs are both regulators and sources of miRNAs, understanding lncRNA–miRNA relationships is important for understanding gene regulation. RNA-seq data from the EMBL-EBI archive were quality-checked with FastQC, aligned with HISAT2 and assembled with StringTie. Transcripts shorter than 200 nucleotides (nt) and other ncRNAs (rRNA, tRNA and snoRNA) were removed, and coding potential was assessed with CPC2 and PLEK, followed by BLASTX against Pfam to remove protein-coding sequences. Differential expression was analysed with NOISeq, and lncRNA–miRNA interactions were predicted with psRNATarget. In total, 7,390 presumed lncRNAs were identified, of which 5,496 were differentially expressed in susceptible and resistant wheat lines. LncRNA targets of 59 miRNAs were analysed; these miRNAs have been reported to regulate plant disease-related genes such as the SPL and NAC families and the cyclophilins TaCYP61, TaCYP62, TaCYP43, TaCYP76 and TaCYP81.

## Introduction

### Triticum aestivum Background

Bread wheat (*Triticum aestivum* L.), known as common wheat, is an annual, predominantly autogamous grass of the family *Poaceae* (tribe *Triticeae*). Its grain supplies a large share of the daily calorific intake in many regions, about 40–50% in Egypt and Turkey and about 20% in the UK. It is also an important source of protein, dietary fibre, B vitamins, minerals and other phytochemicals. Wheat is cultivated in almost all parts of the world, on roughly 225 million hectares each year [1]. It is an allohexaploid (2n = 6x = 42; genome BBAADD) with 21 chromosome pairs organized into three sub-genomes, A, B and D [2]. This young hexaploid species arose only 8,500–9,000 years ago through hybridization between a domesticated free-threshing tetraploid progenitor (BBAA) and *Aegilops tauschii*, the diploid donor of the D sub-genome [1].

Yield losses caused by fungal pathogens are a major threat to global food production. Rusts, smuts, Fusarium head blight, Septoria leaf blotch, tan spot and powdery mildew cause the most serious losses in wheat. Many epidemics on crop plants are triggered by an evolutionary innovation in the pathogen that prevents the host immune system from recognizing it [3].

### Fungal Pathogens and Wheat Rust Interactions

The fungal kingdom is estimated to contain 2.2 to 3.8 million species, traditionally identified by morphological characteristics and by reproductive isolation between novel isolates and known species [3]. Fungi evolve fast and can develop antifungal resistance rapidly. Most fungal pathogens spread systemically from plant to plant and from cell to cell inside the plant, and may reach the tillers without producing any external symptom [4]. Among plant pathogens, fungi are probably the most diverse group of ecologically and economically relevant threats, with outcomes ranging from beneficial interactions to the death of the host.

Fungal spores are dispersed by wind, water or insect vectors and attach to host and non-host surfaces. Secretion of an adhesive extracellular matrix prevents spores from being washed away before penetration [3]. Under favourable conditions the infection cycle starts with spore germination and germ-tube formation, which requires complete cell reprogramming and specific regulatory networks.

Plant pathogens are traditionally classified as biotrophs, hemibiotrophs or necrotrophs according to their lifestyle and mode of feeding. Biotrophs obtain nutrients and energy from living plant cells and survive within the interstitial space of the tissue. Important biotrophic fungi include *Blumeria graminis*, *Puccinia triticina*, *Puccinia graminis*, *Cladosporium fulvum* and *Phytophthora infestans* [3]. Leaf rust, caused by *P. triticina* (Figure 1), is a significant pathogen causing yield losses of 10–20%. It evolves rapidly into new virulent pathotypes that overcome recently deployed resistance (R) genes, despite the discovery of about 100 leaf rust resistance genes (Lr1–Lr100) [5]. The molecular basis of obligate biotroph pathogenicity is not fully understood because of the complex interaction, recognition and signalling with the host.

**Figure 1.**
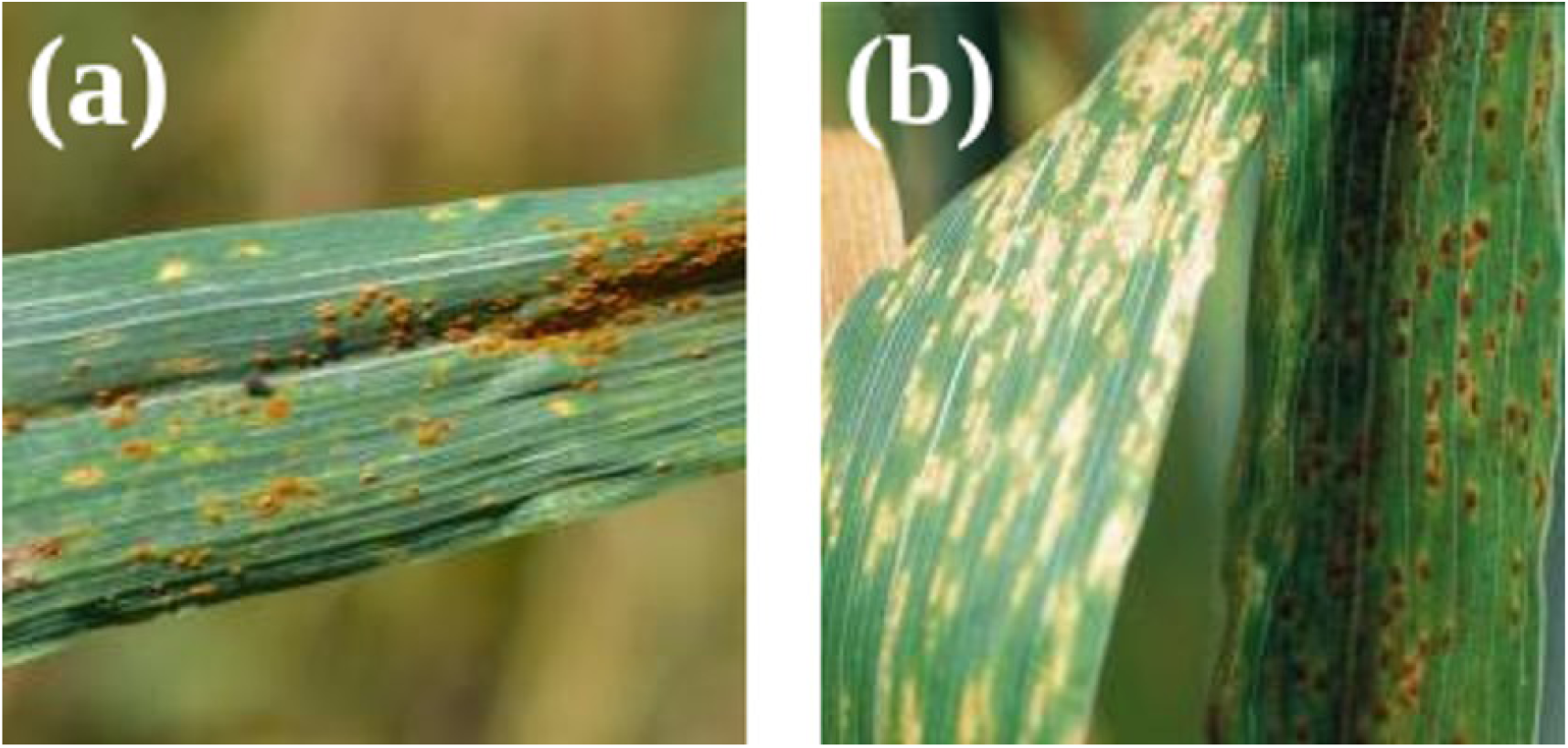
Leaf rust on wheat leaves: (a) rust-coloured spots on the leaf of the Wyalkatchem wheat variety; (b) leaf rust spores, which rarely appear on the back of the leaf except in seedlings.

Hemibiotrophs, such as *Colletotrichum lindemuthianum* and *Magnaporthe* spp., later break down host cell walls with hydrolytic enzymes and feed on the released nutrients [6]. Necrotrophs kill host cells and use their contents for growth; examples include *Alternaria brassicicola*, *Plectosphaerella cucumerina*, *Monilinia* spp., *Sclerotinia sclerotiorum* and *Fusarium* spp. [7].

### Non-Coding RNAs

Non-coding RNAs (ncRNAs) are RNA molecules that do not encode functional proteins. They include conventional housekeeping RNAs, such as transfer RNA (tRNA) and ribosomal RNA (rRNA), and a growing set of regulatory RNAs. ncRNAs differ in size, function and biogenesis, and make up about 80% of the total transcriptome mass of a cell. By size, ncRNAs of 200 nucleotides (nt) or less are small ncRNAs, and those above 200 nt are long ncRNAs [8]. High-throughput sequencing and large-scale transcriptomic analyses have shown that a large proportion of the eukaryotic genome is transcribed into RNAs that do not encode proteins [9].

In addition to protein-coding genes, ncRNAs are increasingly recognized as regulators of plant immunity, and their diverse subcellular localizations may relate to their specific roles in plant cells. Small regulatory ncRNAs include microRNA (miRNA), small interfering RNA (siRNA) and piwi-interacting RNA (piRNA), whereas long ncRNAs include long intergenic ncRNA (lincRNA), natural antisense transcripts (NATs) and circular RNA (circRNA). lncRNAs take part in genome organization, chromatin structure and DNA repair. Many ncRNAs can also be translated into micropeptides, which regulate plant and animal functions, including stress responses.

### Characterization and Biogenesis of lncRNAs

Long non-coding RNAs (lncRNAs) are endogenous single-stranded transcripts longer than 200 nt that do not belong to other well-defined ncRNA classes. Because of their lack of protein coding, poor conservation among species, stage- and cell-type specificity and low abundance, lncRNAs attracted little attention in early years [10]. Their roles in animal growth, development and immunity are well established, whereas studies of lncRNA function in plant immunity lag behind. Many lncRNAs respond to pathogen infection or insect infestation, so biotic stress is a good system in which to study their actions.

According to their position relative to neighbouring protein-coding genes, lncRNAs fall into five groups: sense lncRNAs, on the same strand as the associated gene and overlapping its coding region; antisense lncRNAs, on the opposite strand; intronic lncRNAs, within an intron; bidirectional lncRNAs, on the opposite strand less than 1 kb from the promoter; and lincRNAs, transcribed from the region between two protein-coding genes. In plants, antisense lncRNAs and lincRNAs are the best characterized, and bidirectional lncRNAs are rare.

The biogenesis of lncRNAs is similar to that of mRNAs (Figure 2) [11]. They are usually transcribed by RNA polymerase (Pol) II from intergenic, exonic or distal protein-coding regions, and then undergo 5′ capping, 3′ polyadenylation and sometimes alternative splicing. Some non-polyadenylated lncRNAs appear to be more specific to stress responses. In plants, Pol III, Pol IV and Pol V can also transcribe lncRNAs. Pol III usually produces short, abundant and stable RNAs such as 5S rRNA and tRNA. Pol IV transcripts serve as RDR2 templates for 24-nt small RNAs, whereas Pol V transcripts act as scaffolds that recruit 24-nt small RNAs to complementary genomic loci. Pol IV- and Pol V-derived lncRNAs are poorly characterized because of their low expression and instability.

**Figure 2.**
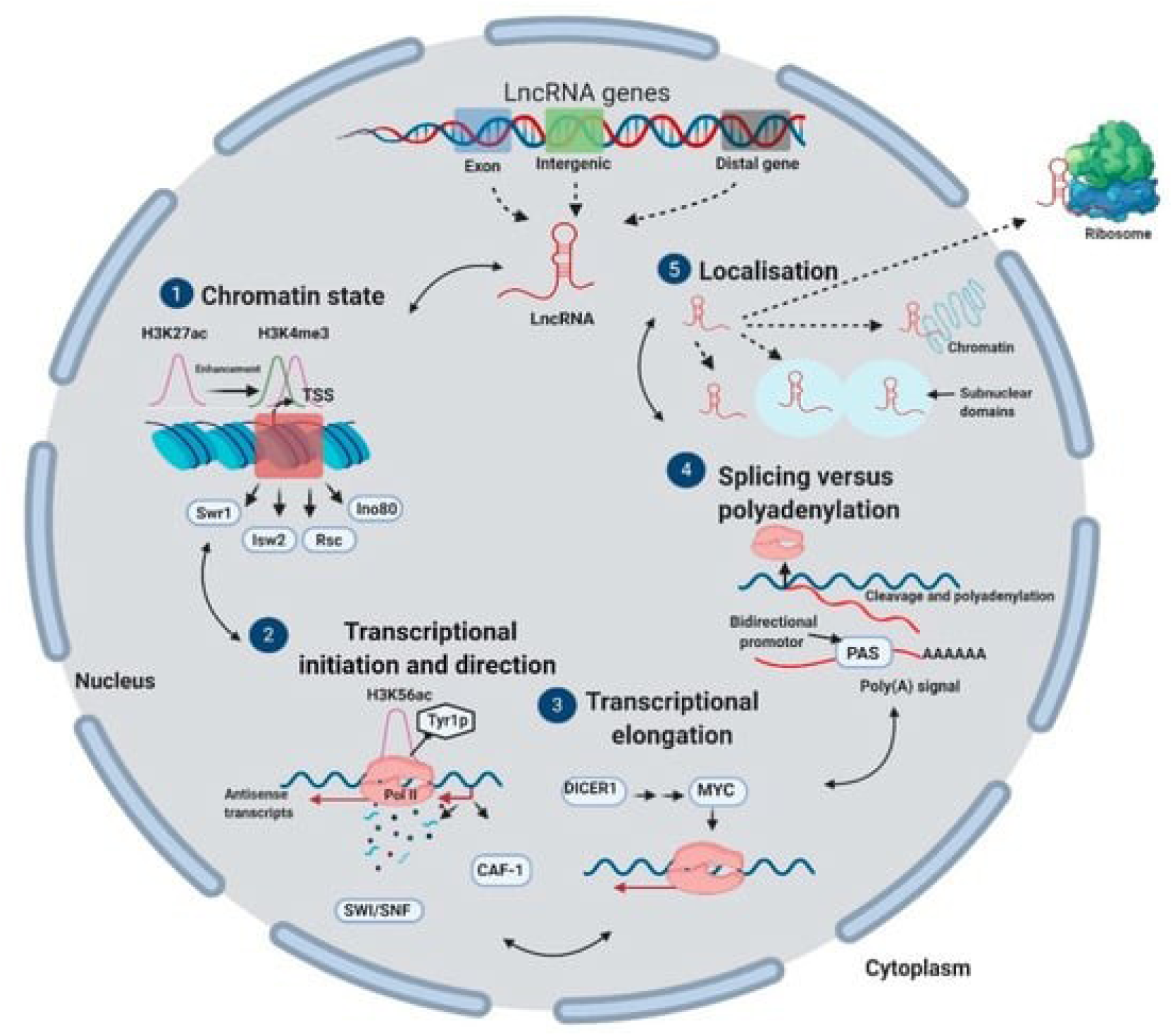
Diagrammatic representation of the biogenesis of lncRNAs.

### Role of lncRNAs in Plant–Pathogen and Wheat Rust Interactions

When wheat plants are attacked by leaf rust, hundreds of downstream genes in signal-transduction pathways show regulated expression. This regulation depends partly on ncRNAs, including miRNAs and lncRNAs. lncRNAs take part in transcriptional and post-transcriptional regulation of gene expression [11]. They may act as precursors of miRNAs or as endogenous target mimics (eTMs) that mimic the real targets of miRNAs and thereby render the corresponding miRNAs ineffective. Leaf rust (*Puccinia triticina*) and powdery mildew (*Blumeria graminis* f. sp. *tritici*) are two important fungal diseases of wheat that cause significant crop damage in epidemic years and threaten food supply [12]. Understanding the resistance mechanism can therefore help to control disease and minimize crop losses.

Leaf rust is generally found on leaves but may also infect glumes and awns. Symptoms begin as small circular to oval yellow spots on the upper leaf surface and develop into orange pustules that may be surrounded by a yellow halo (Figure 1). The pustules release large numbers of spores, which leave an orange dust on the leaf surface, and black spores may appear later, producing mixed orange and black lesions on the same leaf.

Earlier studies identified lncRNAs associated with leaf rust resistance in wheat and showed that specific lncRNAs are upregulated in response to leaf rust infection [13]. Other work showed that certain lncRNAs act as miRNA sponges and thereby regulate defence-related transcripts [14]. Seventy-one wheat lncRNAs were identified in response to powdery mildew; they showed tissue-specific expression patterns, and some functioned in plant immunity as miRNA precursors [15]. However, the roles of lncRNAs in plant–fungus interactions and the underlying mechanisms remain largely unknown.

### Role of Bioinformatics Tools

Identification and characterization of lncRNAs in wheat require a rigorous computational pipeline. FastQC is used for quality control of sequencing data, HISAT2 for genome alignment, SAMtools and BAMtools for alignment file management, and StringTie for transcript assembly. BLASTN verifies sequence homology, PLEK and CPC2 assess coding potential, and BLASTX detects protein-coding sequences. OmicsBox (NOISeq) supports differential expression analysis, and psRNATarget predicts lncRNA–miRNA interactions. In this study, this pipeline was applied to identify lncRNAs in susceptible and resistant wheat lines during leaf rust infection.

## Materials and Methods

The computational pipeline used to identify lncRNAs in wheat–fungal pathogen interactions is shown in Figure 3, and each step is described below.

**Figure 3.**
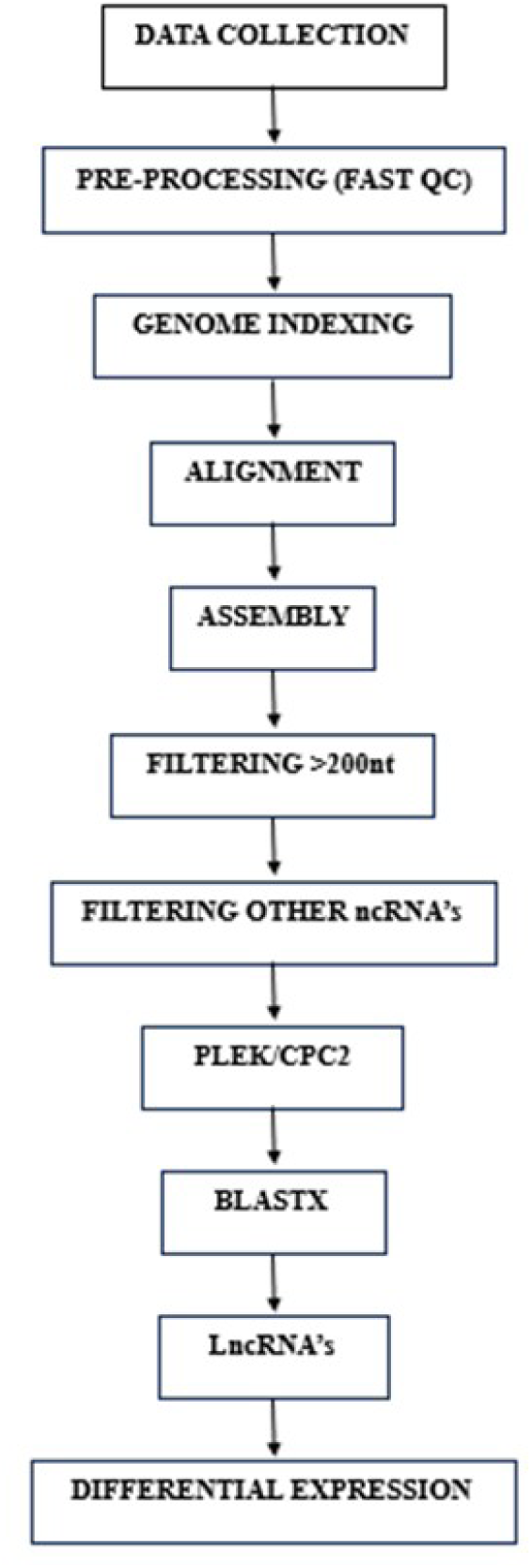
Computational pipeline for the identification of lncRNAs in wheat–fungal pathogen interactions.

### Data Retrieval

RNA-seq data were retrieved from the European Molecular Biology Laboratory–European Bioinformatics Institute (EMBL-EBI) archive using the BioProject and run accession numbers SRR10423617, SRR10423628, SRR10423630 and SRR10423632. These data come from a previous study of Lr28-mediated leaf rust resistance, sequenced on the Illumina NextSeq 500 platform [5]. The four samples are described in Table 1.

**Table 1.** Retrieved data of four different samples (hpi: hours post inoculation)

| **Accession** | **Sample type** | **Label** |
|---|---|---|
| SRR10423617 | Susceptible cultivar HD2329 before infection (0 hpi) | S_0hpi |
| SRR10423628 | Resistant near-isogenic line (NIL) HD2329+Lr28 before infection (0 hpi) | R_0hpi |
| SRR10423630 | Susceptible cultivar HD2329, 96 h after inoculation with the leaf rust pathogen | S_96hpi |
| SRR10423632 | Resistant NIL HD2329+Lr28, 96 h after inoculation with the leaf rust pathogen | R_96hpi |

### Pre-processing

All sequencing data were first processed to eliminate low-quality reads. Quality was checked with FastQC (version 0.11.5), using a Phred score threshold of 30. In the FastQC plots, the y-axis gives the quality score and the x-axis the position in the read. The colour coding of the plots separates high-, medium- and low-quality scores for per-base sequence quality, per-base sequence content, per-sequence GC content, overrepresented sequences, per-base N content, sequence length distribution, adapter content and sequence duplication levels.

### Genome Indexing

Indexing a genome is similar to indexing a book: it is much faster to look up a word in a pre-built index than to read every page. Indices allow the aligner to narrow down the potential origin of a query sequence within the genome, saving both time and memory. HISAT2 is a fast and sensitive alignment program for mapping next-generation sequencing reads (whole-genome, transcriptome and exome data) against a population of genomes as well as against a single reference genome [16].

### Alignment

Alignment was performed with HISAT2 (hierarchical indexing for spliced alignment of transcripts), a highly efficient system for aligning RNA-seq reads, after genome indexing [16]. The wheat reference genome was downloaded from the International Wheat Genome Sequencing Consortium (IWGSC) website (RefSeq v2.1).

### Assembly

Assembly reconstructs the original transcript sequences by merging aligned shorter reads, whereas map-to-reference consists of mapping reads to a reference sequence. Following alignment, transcripts were assembled with StringTie using the IWGSC wheat annotation file as reference.

### Filtering Process

Transcripts shorter than 200 nt were first removed with the SeqKit tool. Other ncRNAs, namely rRNAs, snRNAs and tRNAs, were then removed using the RNAcentral database. BLASTN was performed for homology checking, using the file of ncRNAs longer than 200 nt as the query.

### Coding Potential Calculation

Coding potential was calculated with two tools, CPC2 and PLEK. CPC2 was used for the coding potential calculation of the assembled sequences, and PLEK was used mainly to identify non- coding RNAs. Transcripts common to both CPC2 and PLEK were selected with BLASTN. BLASTX against the Pfam database was then performed to remove protein-coding sequences from the data.

### Differential Expression Analysis

To measure the expression levels of lncRNAs, the matrix of read counts was filtered for lowly expressed lncRNAs using the counts-per-million (CPM) method of the NOISeq package (v2.38.0) before differential expression analysis. Because no replicates were available for any experimental condition, pairwise differential expression analysis was used. NOISeq is a Bioconductor package that can compare samples from two experimental conditions by simulating replicates. The comparisons were S0 vs R0, S0 vs S96, R0 vs R96 and S96 vs R96, and the overlap among them was visualized with Venny 2.1.

### lncRNA–miRNA Interaction Analysis

Interactions between the identified lncRNAs and wheat miRNAs were predicted with the psRNATarget tool, and the lncRNA–miRNA network was visualized in Cytoscape using the source and target nodes obtained from the psRNATarget output.

## Results

### FastQC Outcome

The quality check of the downloaded data produced basic statistics, including per-base sequence quality, sequence length distribution and sequence duplication levels (Figure 4). Based on the FastQC metrics and plots, the samples were of good quality for further processing. The per-base sequence quality plot, one of the key FastQC analyses, shows the distribution of quality scores at each position in the reads, and thereby indicates whether errors occurred during sequencing.

**Figure 4.**
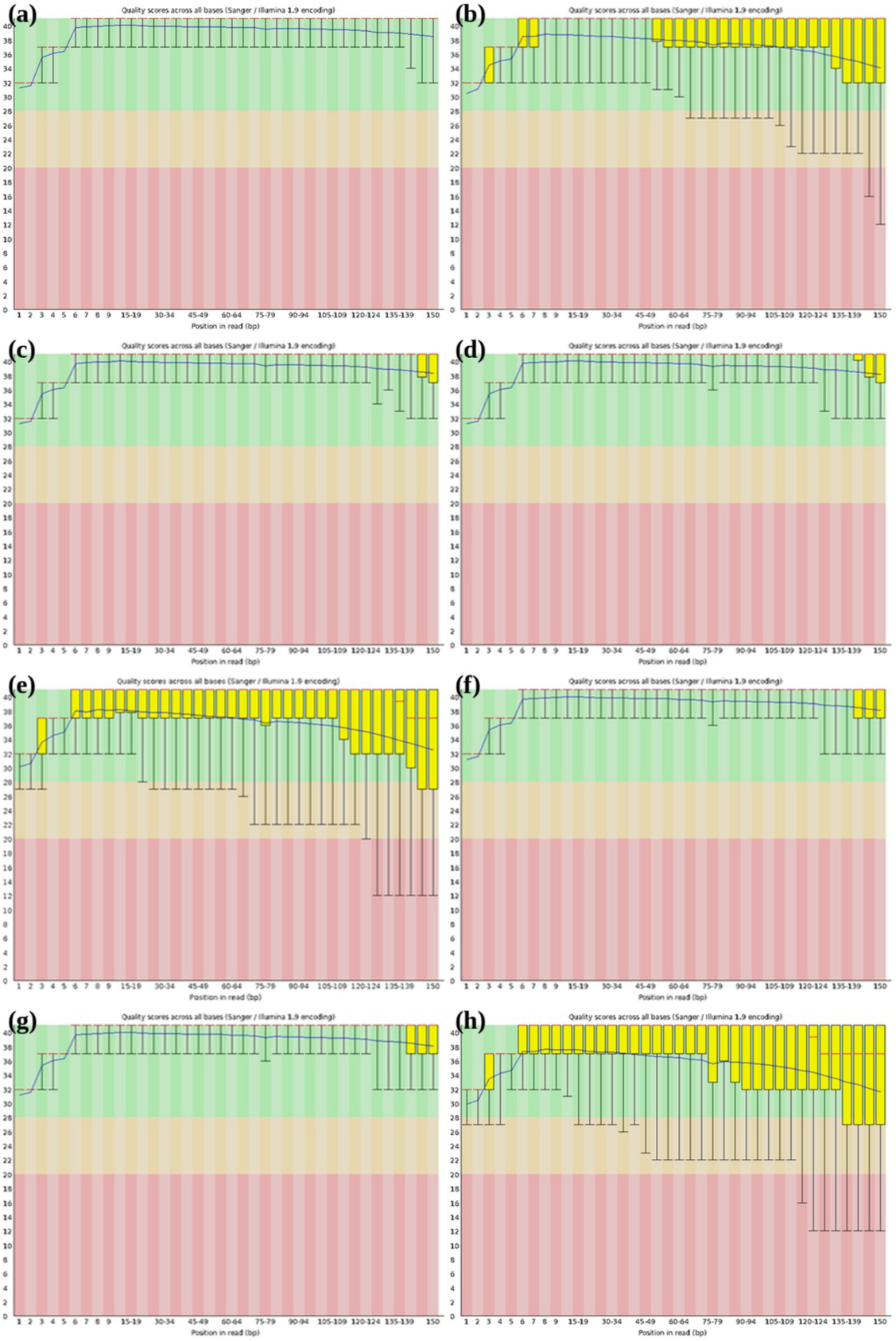
FastQC per-base sequence quality for the collected samples: (a) S0 SRR10423617_1; (b) S0 SRR10423617_2; (c) R0 SRR10423628_1; (d) R0 SRR10423628_2; (e) S96 SRR10423630_1; (f) S96 SRR10423630_2; (g) R96 SRR10423632_1; (h) R96 SRR10423632_2.

### Alignment Process

Alignment was performed with HISAT2. The alignment rate of each data set is given in Table 2; all four samples aligned at more than 89%.

**Table 2.** Alignment rate of the bioproject data ids.

| **Data** | **Sample** | **Alignment rate (%)** |
|---|---|---|
| SRR10423617 | Susceptible (S0) | 91.77 |
| SRR10423628 | Resistant (R0) | 92.61 |
| SRR10423630 | Susceptible (S96) | 90.90 |
| SRR10423632 | Resistant (R96) | 89.60 |

### Identification of Presumed lncRNAs

StringTie assembly generated 205,100 transcripts in total. After removing transcripts shorter than 200 nt, 204,407 transcripts remained. Non-coding RNA databases were downloaded from RNAcentral (https://rnacentral.org/), and rRNA, tRNA and snRNA sequences were removed with BLASTN, leaving 203,865 transcripts. Coding potential calculation then gave 40,855 non-coding transcripts with CPC2 and 79,527 with PLEK. The 38,261 transcripts common to both tools were passed to BLASTX against the Pfam database, which left 7,390 presumed lncRNAs (Figure 5). A total of 5,496 differentially expressed lncRNAs were obtained from the four sample combinations.

**Figure 5.**
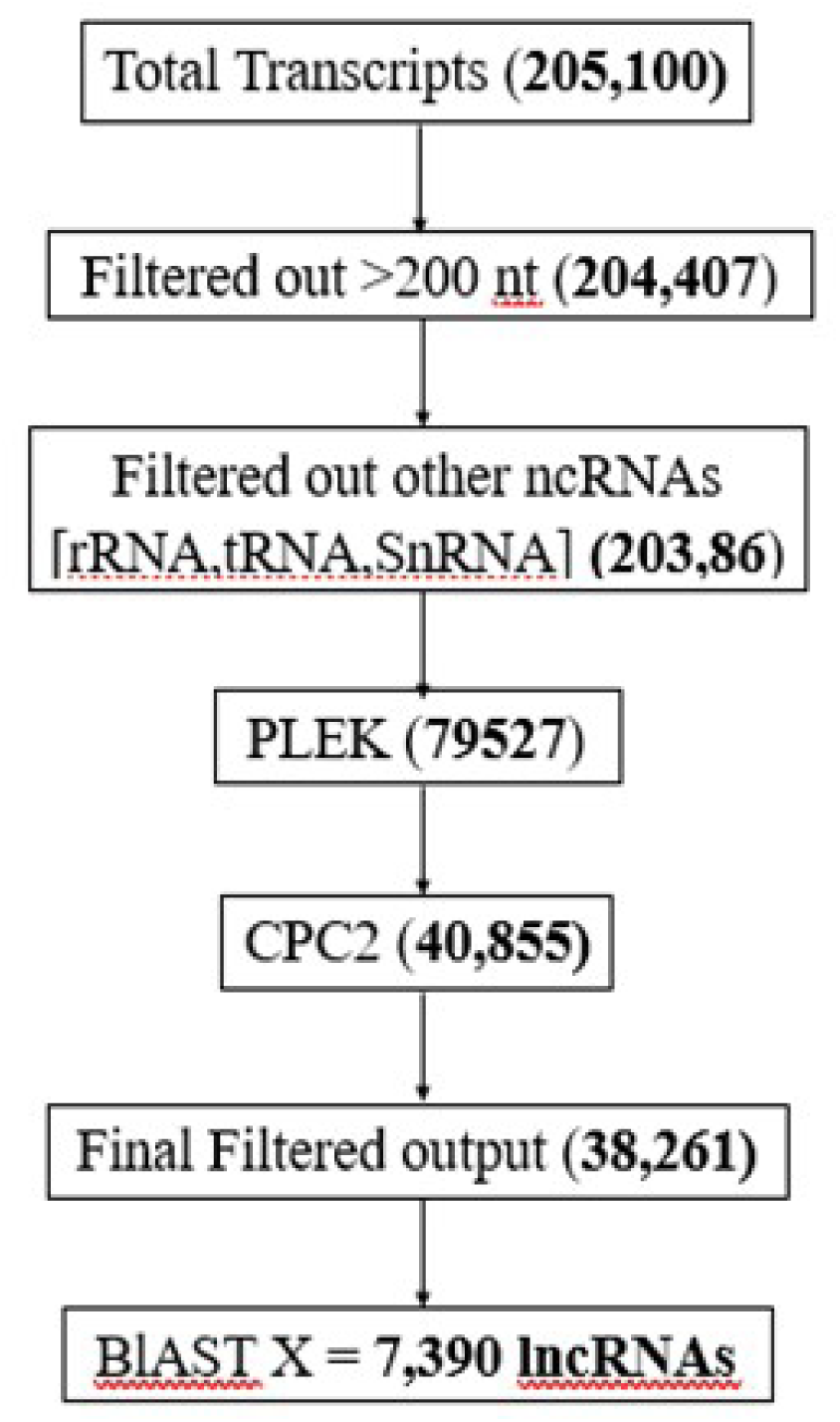
Number of transcripts at each step of the identification of lncRNAs.

### Sub-Genomic Distribution of lncRNAs

Figure 6 shows the distribution of lncRNAs across the A, B and D sub-genomes of wheat: 7,680 transcripts (31.10%) in sub-genome A, 8,868 (35.91%) in sub-genome B and 8,150 (33.00%) in sub-genome D. Previous studies attribute the predominance of lncRNAs in the B sub-genome to a combination of evolutionary history, functional specialization, unique epigenetic regulation and intricate interactions with protein-coding genes [17]. These factors underscore the adaptive complexity of lncRNAs in the B sub-genome, which contributes to the genomic plasticity and stress adaptability of wheat [18].

**Figure 6.**
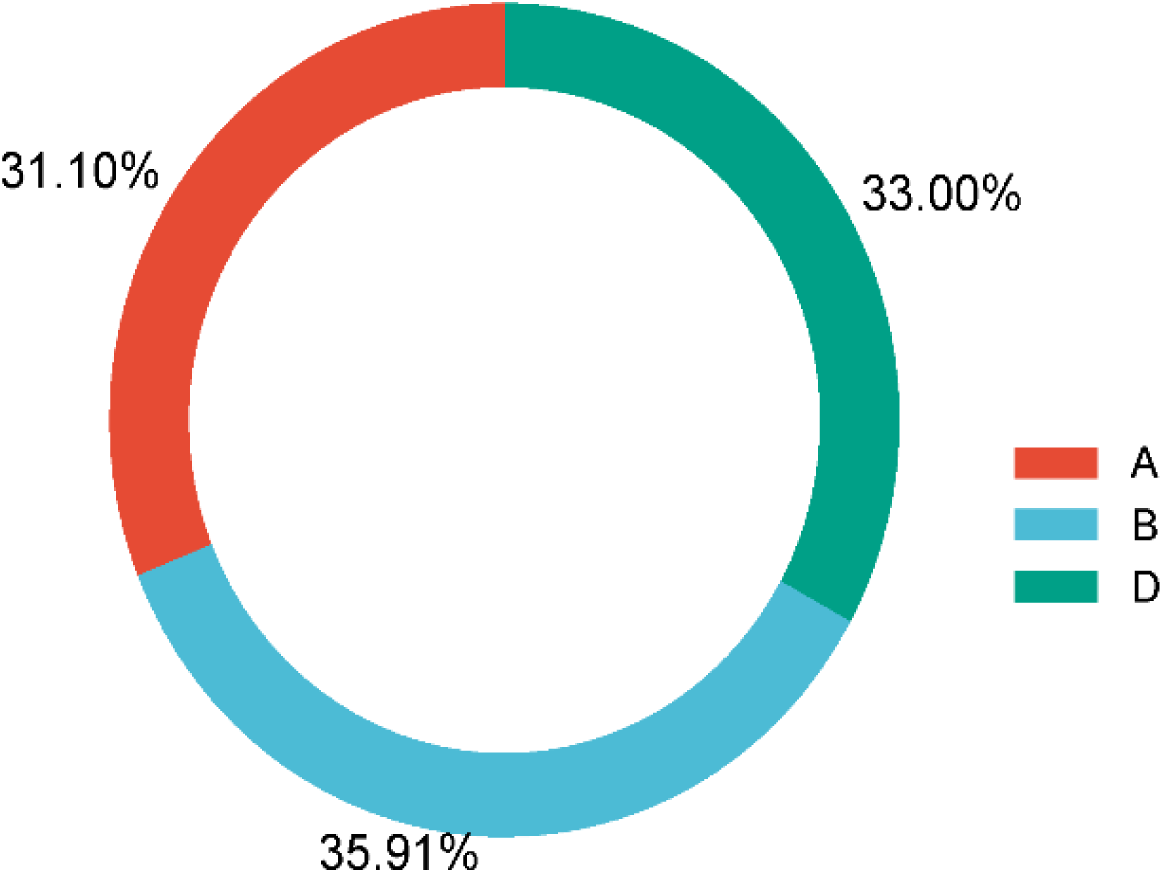
Distribution of lncRNAs across the A, B and D sub-genomes of wheat.

### Distribution of lncRNA Lengths

The length distribution of the lncRNAs obtained from the computational pipeline is shown in Figure 7. Of the 7,390 lncRNAs, 1,281 were 200–300 bp long, 706 were 300–400 bp, 661 were 400–500 bp, 650 were 500–600 bp, 718 were 600–700 bp, 636 were 700–800 bp, 569 were 800– 900 bp, 438 were 900–1,000 bp and 1,731 were longer than 1,000 bp.

**Figure 7.**
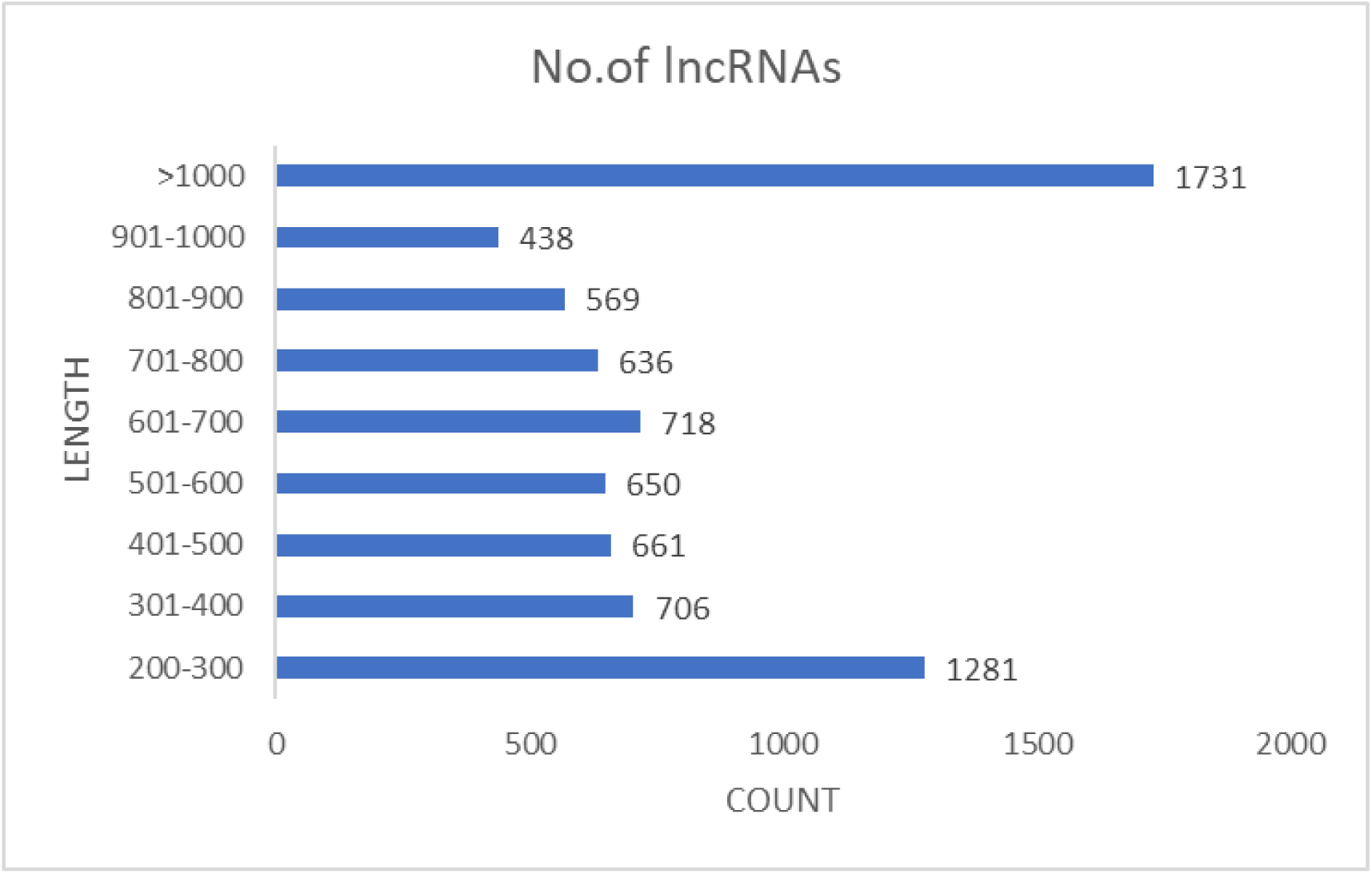
Length distribution of the 7,390 lncRNAs.

### Differential Expression of lncRNAs

The numbers of up- and down-regulated lncRNA transcripts in the four sample combinations (S0 vs R0, S0 vs S96, R0 vs R96 and S96 vs R96) are plotted in Figure 8. In total, 5,496 differentially expressed lncRNAs were obtained with the NOISeq method.

**Figure 8.**
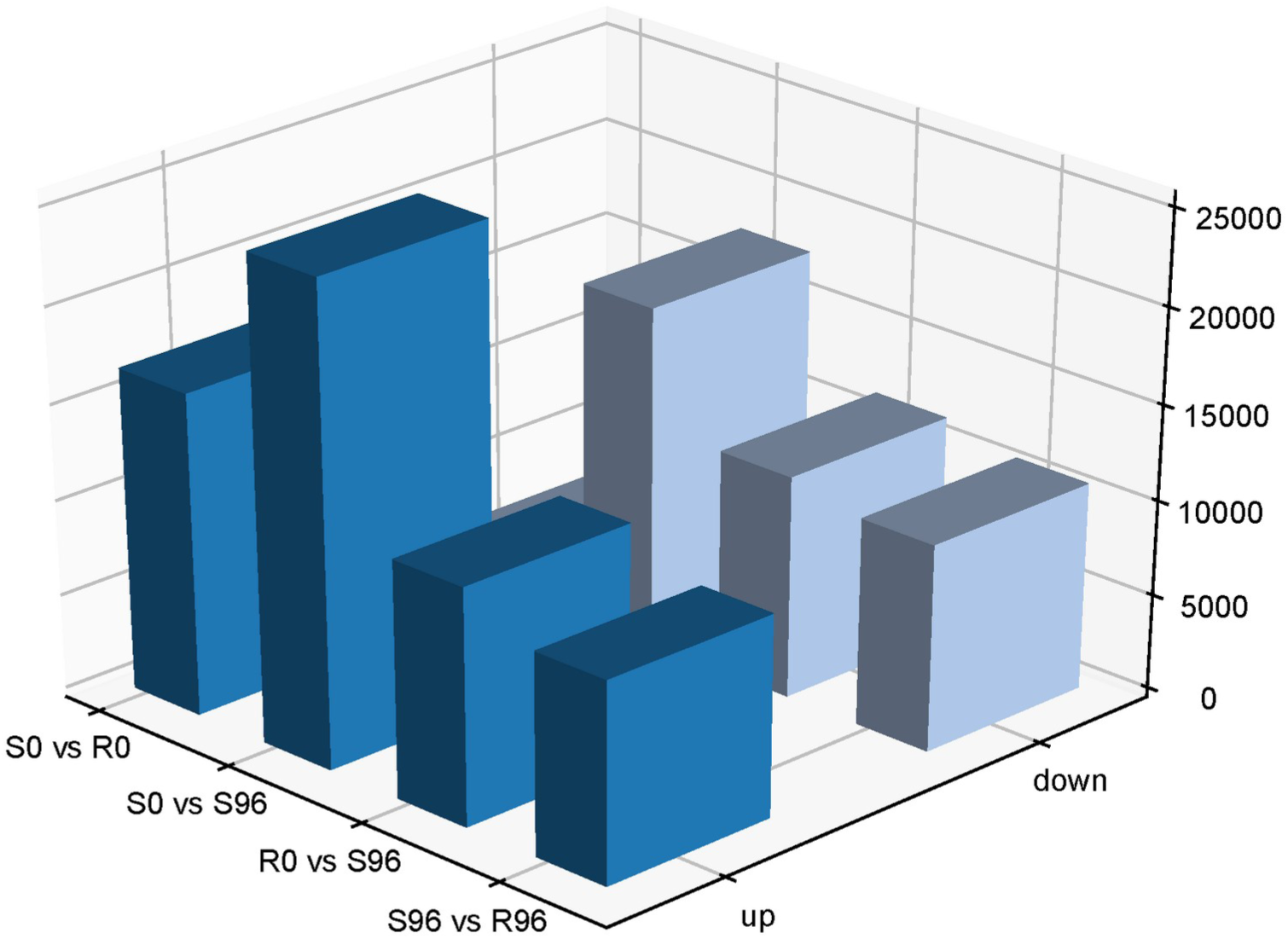
Up- and down-regulated lncRNA transcripts in susceptible and resistant wheat at 0 h and 96 h after inoculation (S0 vs R0, S0 vs S96, R0 vs R96, S96 vs R96).

The Venn diagram in Figure 9 was produced with Venny 2.1 and shows the overlap among the four combinations. A total of 243 lncRNA transcripts were differentially expressed in all four combinations. Some transcripts were exclusive to one combination; for example, 402 differentially expressed transcripts were found only in susceptible plants at 96 h post infection compared with susceptible plants before infection.

**Figure 9.**
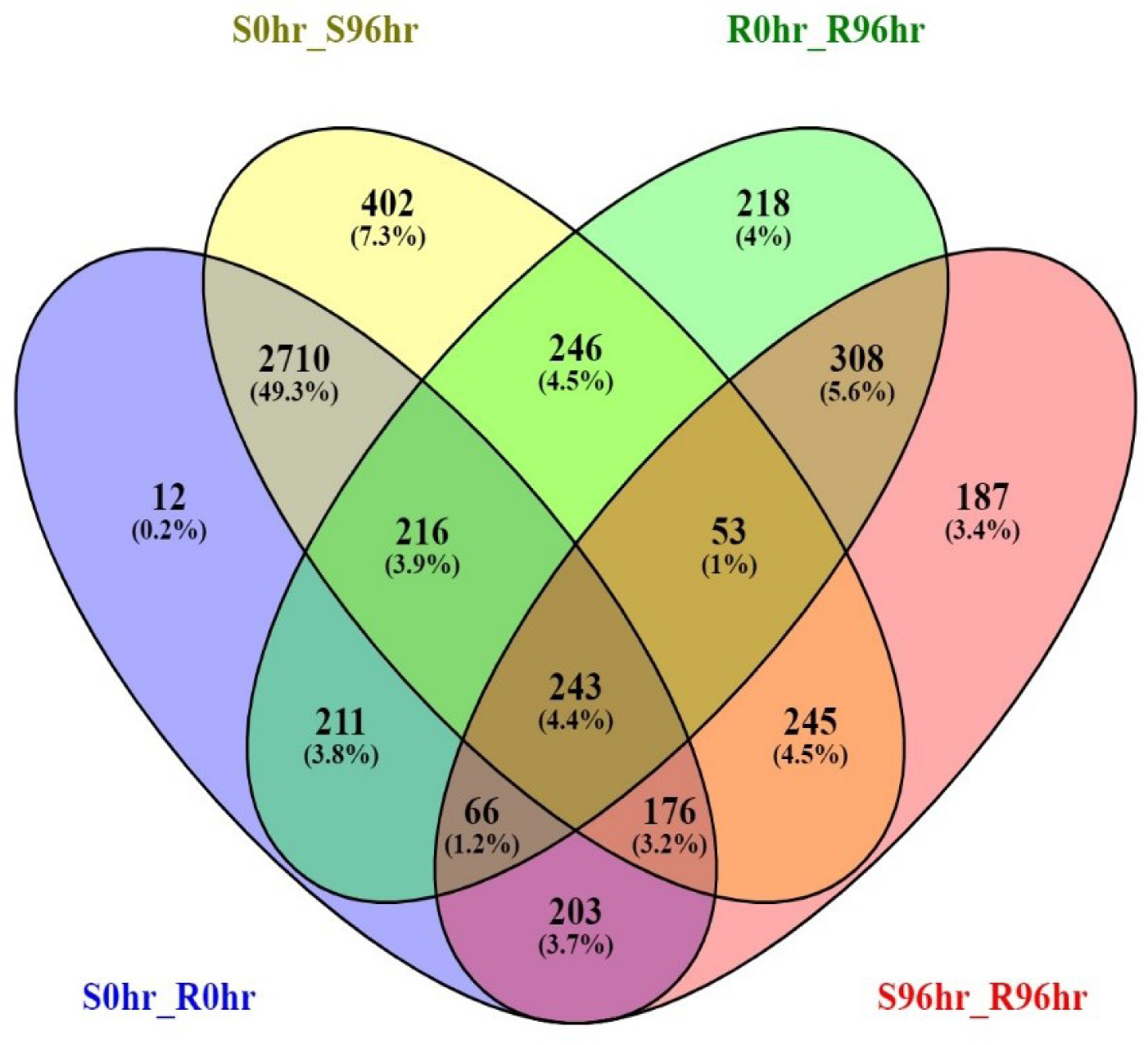
Venn diagram of differentially expressed lncRNAs in susceptible and resistant wheat at 0 h and 96 h.

### Regulatory Network: miRNA Targets in lncRNAs

The miRNA–lncRNA network (Figure 10) was built in Cytoscape from the target lncRNAs obtained with psRNATarget. LncRNA targets of 59 miRNAs were analysed. Table 3 lists representative miRNAs together with their previously reported functions.

**Figure 10.**
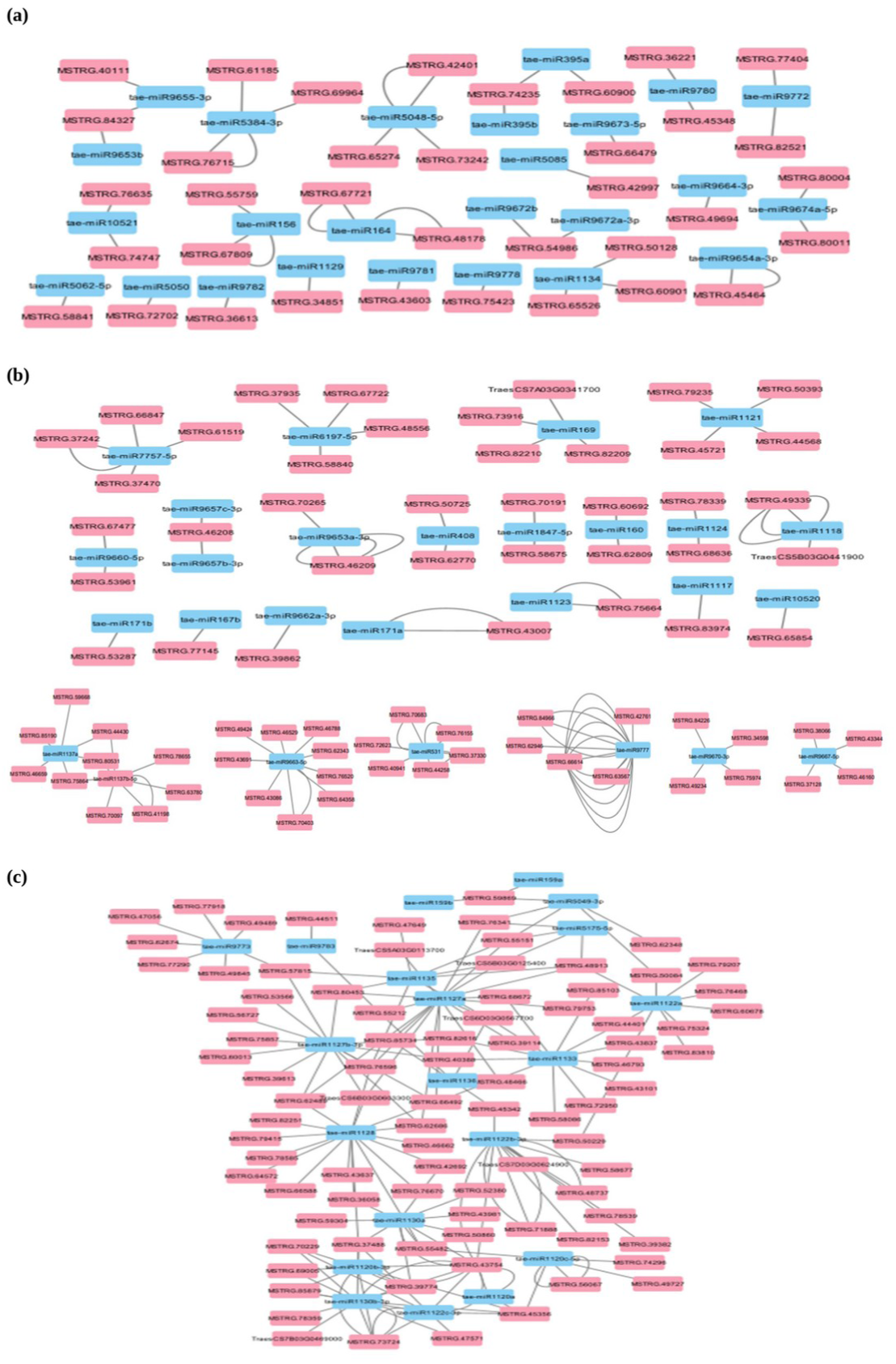
Interaction network between lncRNAs and miRNAs (a)–(c). Pink nodes represent lncRNAs and blue nodes represent miRNAs.

**Table 3.** Lncrna-targeting mirnas and their reported functions.

| miRNA | Reported function |
|---|---|
| tae-miR1120 | Participates in CRY/PHY, auxin and JA signalling pathways [19]. |
| tae-miR9780 | Complementary to Lr34; implicated in resistance of wheat hybrids to *Puccinia triticina*, with Lr34, Lr67 and Lr46 candidate genes [20]. |
| tae-miR1137 | Targets TaCYP24; isomiRs also target anthranilate synthase (tryptophan biosynthesis) [21]. |
| tae-miR5175 | Dehydration-responsive miRNA in *Aegilops tauschii*, the wheat D-genome progenitor, under drought stress [22]. |
| tae-miR1121 | Regulates oxidative reactions under phenanthrene stress in wheat roots [23]. |
| tae-miR1129 | Involved in cellular defence response to infection and in the hypersensitive response [24]. |
| tae-miR1130 | Targets five cyclophilin genes (TaCYP61, TaCYP62, TaCYP43, TaCYP76, TaCYP81) in leaf rust resistance and anthranilate synthase [21]. |
| tae-miR171 | Targets SCARECROW-like transcription factors and regulates root growth in *Arabidopsis* [25]; altered levels change virus symptoms, and over-expression reduces susceptibility [26]. |
| tae-miR156 | Acts through SPL transcripts to lower miR172; reduced in crown tissue of dwarf lines at normal temperature [27]. |
| tae-miR408 | Target TaCLP1 enhances tolerance to high salinity, copper stress and stripe rust [7]. |
| tae-miR395 | Targets a WRKY transcription factor; many PR-protein genes in apple leaf spot resistance [28]. |
| tae-miR164 | Target is a NAC transcription factor (NAM subfamily) that regulates wheat resistance to stripe rust [29]. |
| tae-miR169 | Together with methylation, regulates disease resistance genes in wheat [30]. |
| tae-miR167 | With other miRNAs, modulates defence against stem rust through R-gene-independent pathways [24]. |
| tae-miR160 | Regulates defence against anthracnose through ARF10 and TIR1 targets [31]. |
| tae-miR159 | Derived vsiRNAs regulate *taMyb3* and enhance resistance to *Puccinia striiformis* [32]. |
| tae-miR5050 | Fe-responsive miRNA with tissue-specific regulatory role during Fe deficiency [33]. |
| tae-miR5049 | Predicted roles in Wnt signalling and drought response [34]. |
| tae-miR5384 | Expression changes after *P. triticina* inoculation; possible targets in unsaturated fatty acid formation; increases after chitosan treatment [20]. |
| tae-miR1127 | Wheat small RNA response to an obligate pathogen, shaped by domestication of high-copy transposons [35]. |
| tae-miR9663 | Potential biomarker for screening wheat varieties with low cadmium accumulation [36]. |
| tae-miR531 | Associated with potassium-deficiency tolerance and phenanthrene metabolism in wheat [37]. |
| tae-miR5048 | Enhances salt adaptation in autopolyploid *Hordeum bulbosum* compared with its diploid ancestor [38]. |
| tae-miR1118 | Targets expansins and aspartyl-protease proteins; modulates Fe redox status in hexaploid wheat [33]. |

## Discussion

In recent years, lncRNAs have emerged as major regulators of genes involved in biotic and abiotic stress responses in plants [11]. In wheat, few lncRNAs have been identified that play a role in abiotic stress responses, seed germination and disease resistance. Although progress has been made towards understanding the molecular mechanisms behind wheat resistance to fungi, little is known about the potential roles of lncRNAs in the response to fungal pathogens [39]. Leaf rust is one of the most destructive fungal diseases of wheat, reducing grain quality and causing large economic losses, and many molecular events are activated to reprogram plants to resist pathogens during plant–pathogen interaction.

In this study, RNA-seq data from susceptible and resistant wheat infected with leaf rust were retrieved and quality-checked, with Phred scores above 30. HISAT2 alignment rates exceeded 89% for all four samples, indicating that the majority of reads could be aligned to the reference genome. Assembly and downstream filtering showed that lncRNAs were most abundant in the B sub-genome, which earlier studies attribute to evolutionary history, functional specialization and unique epigenetic regulation [17], [18]. The lncRNA lengths clustered mainly in two classes: 1,731 transcripts longer than 1,000 nt and 1,281 transcripts of 200–300 nt. In total, 5,496 differentially expressed transcripts were obtained, and 243 were common to all four combinations (Figure 9) [40].

Compared with previous studies, the commonality of differentially expressed lncRNAs in both susceptible and resistant lines suggests conserved roles in basic immune responses and cellular processes [41]. These lncRNAs might be involved in general defence mechanisms, such as oxidative stress response, hormone signalling and cell wall modification, that are triggered in both lines regardless of resistance status [42].

The results also map interactions between miRNAs and lncRNAs in the regulation of target genes [43]. As summarized in Table 3, many of the miRNAs that interact with the identified lncRNAs have known roles in plant disease and stress responses. For example, miR1130 targets cyclophilin genes in leaf rust resistance, miR9780 is complementary to Lr34, miR164 regulates a NAC transcription factor involved in stripe rust resistance, and miR159 and miR167 modulate defence responses to stripe rust and stem rust. Similar lncRNA–miRNA regulatory associations have been described for *Puccinia striiformis* in wheat [44], in grapevine [45] and in wheat inoculated with *Rhizoctonia cerealis* [46]. Taken together, these findings indicate that the identified lncRNAs interact with miRNAs that have established roles in plant defence against fungal pathogens.

## Conclusion

Regulatory RNAs such as lncRNAs play important roles in biological processes and metabolic activities in many plants through gene regulation, but their study in wheat under biotic stress is very limited. This study identified lncRNAs and their expression in response to a fungal pathogen in wheat. The computational analysis identified 7,390 lncRNAs in *T. aestivum*, of which 5,496 were differentially expressed. The lncRNAs interact with 59 wheat miRNAs that target many genes involved in defence against fungal pathogens. The results provide useful information for further exploration of the activity of long non-protein-coding genes in defence against fungal pathogens in wheat, and an understanding of the mechanism of gene regulation will contribute to breeding programs for resistant wheat.

## Acknowledgements

B. Yaswitha thanks Dr. V. B. Sameer Kumar, Head, Department of Genomic Science, for providing departmental facilities, and Dr. Padmesh Pillai, Dr. M. Nagarajan and Dr. Tony Grace for their support and cooperation. B. Yaswitha also thanks Ms. Gayatri K and Ms. Nikitha for their support, the non-teaching staff of the Department of Genomic Science for administrative and laboratory facilities, and the Central University of Kerala for providing an ideal academic environment.

## Data availability

The RNA-seq data analysed in this study are publicly available from the EMBL-EBI/NCBI sequence read archives under run accession numbers SRR10423617, SRR10423628, SRR10423630 and SRR10423632.

